# The effect of wrist posture on extrinsic finger muscle activity during single joint movements

**DOI:** 10.1101/769414

**Authors:** Carl R. Beringer, Misagh Mansouri, Lee E. Fisher, Jennifer L. Collinger, Michael C. Munin, Michael L. Boninger, Robert A. Gaunt

## Abstract

Wrist posture impacts the muscle lengths and moment arms of the extrinsic finger muscles that cross the wrist. As a result, the electromyographic (EMG) activity associated with digit movement at different wrist postures may also change. Therefore, we sought to quantify the posture-dependence of extrinsic finger muscle activity. Fine-wire bipolar electrodes were inserted in the extrinsic hand muscles of able-bodied subjects to record EMG activity during wrist and finger movements in various postures. EMG activity of all the recorded finger muscles were significantly different (p<.05, ANOVA) when performing the same movement in five different wrist postures. EMG activity varied by up to 70%, with the highest levels of activity observed in finger extensors when the wrist was extended. Similarly, finger flexors were most active when the wrist was flexed. For the finger flexors, EMG variations with wrist posture were most prominent for index finger muscles, while the EMG activity of all finger extensor muscles were similarly modulated. The extrinsic finger muscles also showed significant activity during wrist movements with the digits held still regardless of finger posture, suggesting that they may play a role in generating torque during wrist movements. Finally, we developed a pair of generalized classifiers that show that finger muscle EMG can be used to predict wrist posture. These results may impact the design of biomimetic control algorithms for myoelectric prosthetic hands, but further work in transradial amputees is necessary to determine whether this phenomenon persists after amputation.

## Introduction

The human hand is a complex biomechanical system which relies on intrinsic active and passive forces to maintain postures and perform movements. The movement of a single finger alone generates torque at the wrist^1^ that must be counteracted to maintain wrist posture. Single digit motion also generates torque in neighboring digits^2,3^, a phenomenon known as force enslavement, which must also be counteracted by other finger muscles. The force enslavement effect has been suggested to be a combination of passive mechanical coupling driven by tendinous connections between digits as well as active coupling from higher order motor commands^2,4^. While there has been an effort to characterize the resulting electromyography (EMG) activity in these scenarios, prior work has primarily used surface EMG recordings and/or examined only a single muscle at a time^5–8^, leaving an incomplete understanding of how the many extrinsic hand muscles coordinate for single and multiple digit movements. Intramuscular EMG is an alternative method of recording EMG signals by placing fine-wire electrodes through the skin directly into muscles. Intramuscular recordings have been used for over 50 years to characterize muscle activity^9–11^ and provides focal recording from a single target muscle that is difficult or impossible to achieve with standard surface EMG. Fine-wire electrodes can also record distinct signals from deep muscles and the different compartments of the multicompartment finger flexor and extensor muscles^12–15^.

Extensive work has demonstrated that coupling between the digits is composed of mechanical^16,17^ and neural components^2,14,15,18,19^, although the influence of inter-digit coupling on the coordination of single and multiple degrees of freedom (DOF) movements is not well understood. Experiments in humans have shown that electrical stimulation induces more focal and independent finger forces compared to volitional movement, suggesting that neural coupling does in fact result from cortical motor control strategies^14^. The influence of coupling is also visible during EMG recordings; an example may be seen in the work by Leijnse *et al*.^5^, which shows phasic EMG activity in the extensor digiti minimi during a thumb tapping task. However, EMG activity of other muscles which appear to be functionally unrelated to the actuation of a joint may contain unique information that is relevant for prosthetic control. Improved EMG decoder performance was observed by Smith *et al*.^20^ when all recorded extrinsic hand muscles were included in the decoder regression equations for each DOF.

Wrist posture is known to have mechanical effects on the fingers, the most direct being motion of the relaxed fingers while the wrist is flexed and extended. For example, when the wrist is extended the fingers naturally curl into a tenodesis grasp^21^. This occurs because the extrinsic finger flexor muscles span the wrist and are subject to lengthening based on wrist angle, which in turn causes an increase in passive tension in the flexor muscles^22^. The resulting increase in force has also been shown to affect the EMG activity necessary to maintain the static pose of a digit^23^. Grip force is also affected by wrist posture, which may be observed through alterations in EMG activity^24,25^. Duque *et al*.^24^ quantified this relationship between grip force and EMG as a series of nonlinear models for flexed, extended, and neutral postures, demonstrating that wrist posture alters the relationship between grip force generation and EMG, but it is unknown whether the posture/EMG relationship is effected in low force movements. In addition, a further motivation to study the relationship between extrinsic finger muscle EMG activity and wrist posture is the potential impact on the design of biomimetic-inspired musculoskeletal model-based control systems for prosthetic hands. Dexterous prosthetic hands exist, but a significant barrier to adoption is a lack of control algorithms which may take advantage of the high degree-of-freedom control that is offered^26^. Biomimetically-inspired control algorithms could possibly improve control^27,28^, thus we sought to improve the understanding of the relationship between extrinsic finger muscle EMG activity and wrist posture.

In this study we used intramuscular EMG to target the compartments of the flexor digitorum profundus (FDP), flexor digitorum superficialis (FDS) and extensor digitorum (ED), as well as the extensor digiti minimi (EDM) and the extensor indicis proprius (EIP) muscle, and sought to characterize the effect of changing wrist joint angles on EMG activity of the extrinsic finger muscles during structured hand movements. We also examined how the extrinsic finger muscles may assist in wrist movements themselves.

## Methods

### Experimental Summary

11 healthy, able-bodied subjects (7 males, 4 females) were included in this study. Subject ages ranged from 25-32 with an average age of 28.2±2.7 years. All subjects provided informed consent prior to any experimental procedure and all procedures were approved by the University of Pittsburgh and Army Research Labs Institutional Review Boards. Sixteen percutaneous EMG electrodes were implanted in the extrinsic hand muscles that control the major finger and wrist flexion and extension movements. Subjects wore a glove fitted with electromagnetic trackers to capture hand kinematics. Subjects then performed an extensive series of single joint movements with varying finger and wrist postures while EMG and kinematics were recorded.

### Percutaneous EMG Recording

Percutaneous fine wire bipolar electrodes (30 and 50 mm long paired fine wire needle electrodes, Chalgren Enterprises, Inc.) were placed in sixteen of the following muscle locations and compartments under ultrasound guidance by an experienced physician: the 2^nd^ and 4^th^ compartments of the extensor digitorum (ED2, ED4), the extensor digiti minimi (EDM), flexor digitorum superficialis (FDS2-4), flexor digitorum profundus (FDP2-5), flexor pollicis longus (FPL), abductor pollicis longus (APL), extensor pollicis longus (EPL), extensor indicis proprius (EIP), flexor carpi ulnaris (FCU), flexor carpi radialis (FCR), extensor carpi radialis longus (ECRL), extensor carpi ulnaris (ECU), pronator teres (PT), and supinator (SUP) muscles. The 2^nd^ and 4^th^ compartments of the extensor digitorum were selected for the greatest separation between signals. We broadly targeted the extrinsic hand muscles during implantation to achieve a representation of the extrinsic hand muscles which controlled the major degrees of freedom of the hand. Muscle locations were identified using palpation and ultrasonographic visualization techniques during instructed movements to guide electrode insertion^13^. Intramuscular EMG recordings were digitized with a multichannel neural recording system (Grapevine Neural Interface System with Surf S2 headstage, Ripple, Inc) at 30 kHz.

### Kinematic Motion Tracking

Hand and arm kinematics were recorded using an electromagnetic tracking system (trakStar, Ascension Technology, Inc.) integrated into the MotionMonitor (Innovative Sports Training Inc., http://www.TheMotionMonitor.com) recording software. The tracking sensors were attached to a glove, and located over the proximal, intermediate, and distal phalanges of the index finger; the proximal and distal phalanges of the middle, ring, and little finger; the metacarpal and phalanges of the thumb; and the dorsal center of metacarpals (**Fig. 1a**). Tracking sensors were also placed over the distal portion of the radius and lateral aspect of the biceps to track the position of the arm. Subject-specific arm and hand segments were then created using the digitization process in the MotionMonitor software. The OpenSim v3.3 Inverse Kinematics tool^29^ was then used to extract the joint angles from the .trc files using a scaled musculoskeletal model^30^.

**Figure 1.**
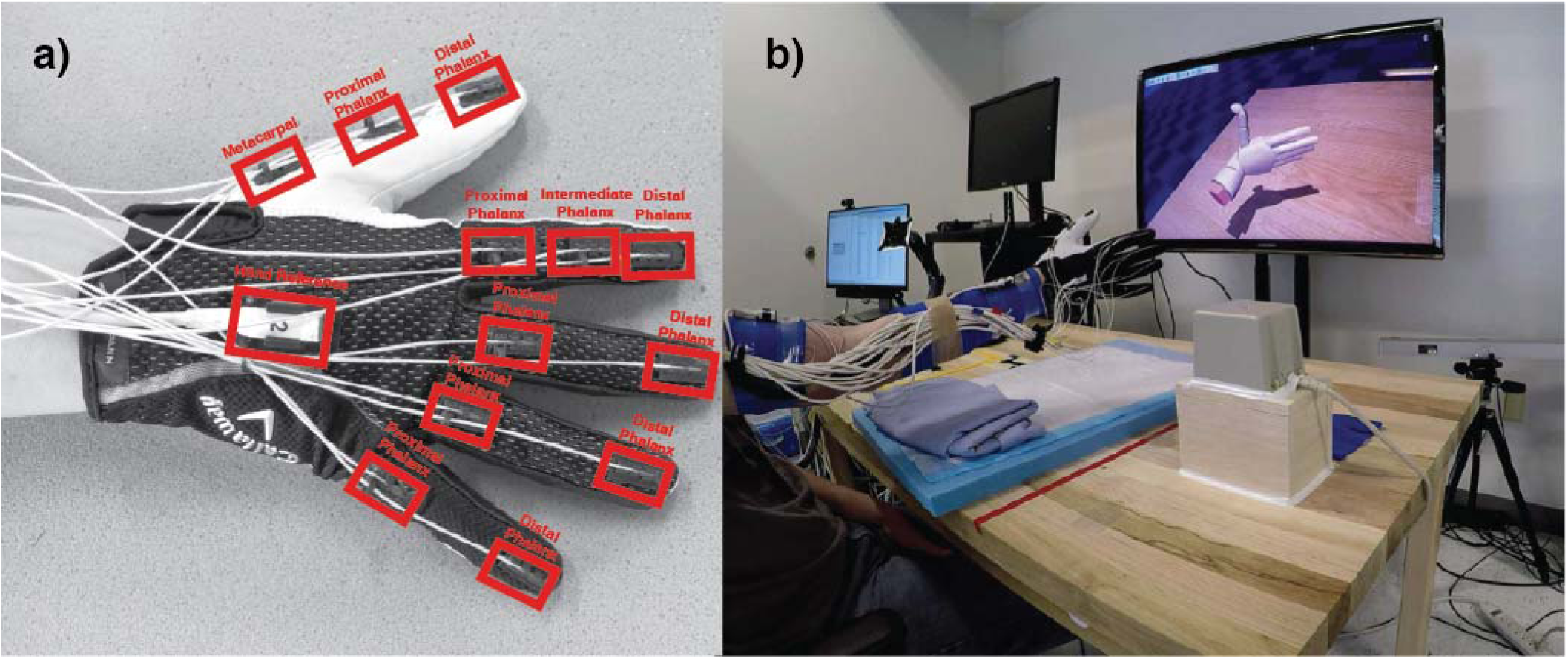
Overview of the experimental setup. **a)** Electromagnetic tracking glove that subjects wore during the experiments. The red rectangles show placement of electromagnetic sensors in relation to the joints of the fingers. **b)** Subjects viewed and followed a video which demonstrated the hand posture, movement, and timing.

### Experimental Tasks

Subjects were asked to follow the movements of a virtual hand displayed on a computer screen (**Fig. 1b)**. Each movement exercised a single DOF of the fingers or wrist for ten repetitions at 1 Hz. D2-D5 flexion/extension and wrist flexion/extension movements were tested. For D2-D5 movements, subjects were asked to produce coupled motions of the metacarpal phalangeal joint along with the proximal and distal interphalangeal joints, with a focus on moving the metacarpal phalangeal joint through 90 degrees. All fingers motions were repeated while the subject maintained the wrist in a neutral posture (hand aligned with the forearm and thumb pointing up), or with the wrist in a flexed, extended, pronated, or supinated position at the limit of the wrist range of motion. During wrist movements, the fingers were held in either flexed or extended positions and the exercises were performed while the wrist was held neutral, pronated, or supinated. Pronation/supination movements were only performed during neutral wrist postures. The initial posture for D2-D5 movements was with all fingers extended and the wrist in one of five positions. For wrist movements, the initial starting pose was with the wrist held neutral and the fingers either flexed or extended, depending on the trial. A list of all performed trials is provided in **Supplementary Table S1**.

### Data Processing

Kinematic and EMG signals were processed offline using Matlab (The Mathworks, version 2016b). Kinematic data were smoothed with a 4^th^ order low-pass Butterworth filter at 10 Hz to remove noise. Single-ended intramuscular EMG recordings were differenced, then high-pass filtered with a 4th order high-pass Butterworth filter at 10 Hz before undergoing a pre-processing step (see below) to remove electromagnetic noise generated by the kinematic motion tracking system **(Fig 2a)**. Following this, EMG data were band-pass filtered with a 4^th^ order Butterworth between 100 to 4000 Hz to remove movement artifacts, line noise, and high frequency noise. Potential extrinsic finger muscle recordings were excluded during the data analysis phase if the EMG activity from an electrode was not modulated by single digit movement, or if the electrode recordings were significantly contaminated by artifacts due to the fragility of the fine-wire electrodes. Artifacts were identified based on signal amplitude and shape. Events with peak voltages much greater than those occurring during a maximal voluntary contraction, or events with large instantaneous changes in amplitude or that did not resemble typical EMG activity or motor unit action potentials were considered artifacts. If multiple electrodes were in the same muscle or muscle compartment for a subject, we included only a single electrode with the highest signal-to-noise ratio.

**Figure 2.**
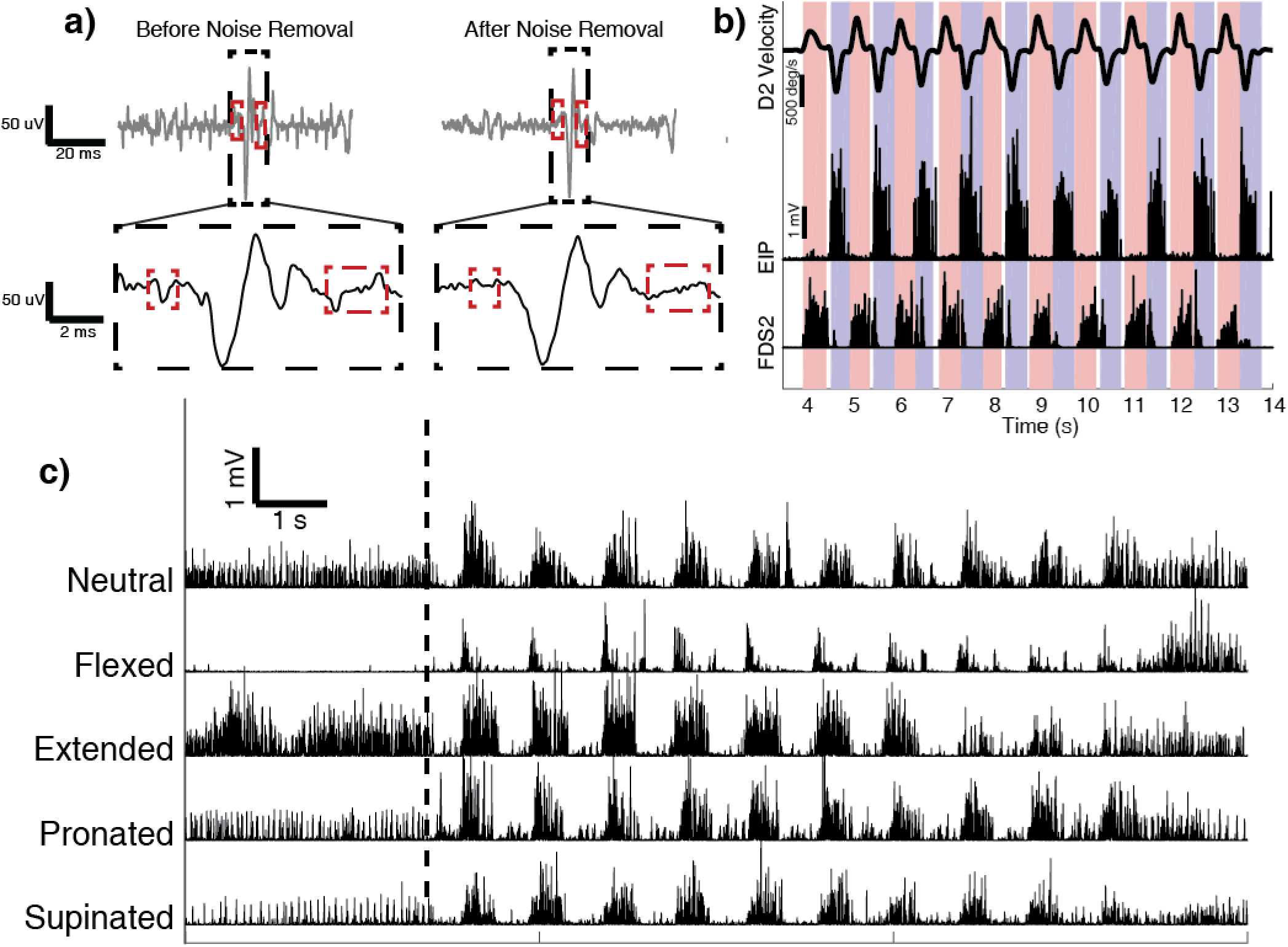
Overview of data processing and effects of wrist posture on EMG activity **a)** Comparison of EMG activity during rest before (left) and after (right) electromagnetic noise removal. Blocked in red are examples of electromagnetic artifacts introduced by the kinematic tracking system and the signal after removal. Left: EMG activity after high pass filtering to remove DC offset and motion artifacts. The artifacts may be observed boxed in red. Right: The same EMG activity after electromagnetic noise removal. A scaled signal has been subtracted from the signal while preserving the EMG activity present during the same time period. **b)** D2 MCP joint velocity and EMG activity of EIP and FDS2 during repeated flexion (red) and extension (blue) movements. The unshaded region between movements represents brief holding periods that were not included in calculations of integrated EMG activity. **c)** Rectified and processed EMG activity of ED4 during repetitions of D4 flexion and extension in neutral, flexed, extended, pronated, and supinated postures. The dashed vertical line shows the task start cue. The hand was held in these static postures for approximately 4 seconds before performing 10 movement repetitions.

The Ascension trakSTAR motion tracking system provides the ability to track the location of a sensor in space by creating a pulsed electromagnetic field in the workspace. This enables sensors to be tracked even when obscured. However, the changing electromagnetic field was also captured by the EMG recording system as a series of pulses occurring at the pulse frequency of 100 Hz. We removed this noise from recordings offline while preserving the EMG signals by generating a template pulse train, and then subtracting it from the data. An example of the EMG signal before and after electromagnetic noise removal is shown in **Fig. 2a**.

### Data Analysis and Statistics

We divided the EMG data for every trial into flexion and extension phases using the kinematic recordings **(Fig. 2b)**. The angular velocities were low-pass filtered with a zero-phase digital filter at 10 Hz. We detected the movement phases by first identifying the positive and negative peak velocities. Movement onset was then detected based on the first zero velocity point preceding each peak, while the end of a movement was the first zero velocity point after a peak. The filtered EMG data were integrated over the relevant movement windows and divided by the duration of the movement to quantify the EMG activity over the period. For example, the flexor digitorum superficialis was integrated during flexion movements and the extensor digitorum was integrated during extension movements.

#### Finger Movements

Integrated EMG activity for single digit movements were grouped across subjects based on the muscle in which electrodes were placed. For the muscles with multiple compartments (ED, FDS, and FDP), integrated EMG data were separated based on the compartment and digit actuated. A list of each muscle analyzed and the number of subjects per muscle are shown in **Supplementary Table S2**. The mean integrated EMG activity for each muscle and subject was calculated for the five postures: neutral, flexed, extended, pronated, and supinated **(Fig. 2c)**. The integrated EMG data were then normalized to the highest of the means. The normalized integrated EMG activity of each muscle and compartment were tested for significant differences using a two-way ANOVA with subject and wrist posture (neutral, flexed, extended, pronated, and supinated) as factors. Post-hoc pairwise testing to identify differences in EMG activity for movements in the postures assessed was performed using Tukey’s honest significant differences (HSD) test for multiple comparisons.

#### Wrist Movements

For each subject and electrode, the mean integrated EMG activity for each finger muscle was calculated for the four combinations of finger postures and wrist movements: fingers extended (‘flat’) during wrist flexion and extension and fingers flexed (‘fist’) during wrist flexion and extension. For the four movements, the EMG activity of each muscle and subject was normalized to the maximum activity to compare across subjects. The normalized EMG activity of ED, EDM and EIP muscles were combined into a single ‘extensors’ group, while FDS and FDP were combined into a ‘flexors’ group. Since the data were not normally distributed (p<.05, Kolmogorov-Smirnov test,) a Kruskal-Wallis test was conducted on the EMG activity of the extrinsic finger flexors and extensors to detect whether EMG activity of finger flexors and extensors differed based on finger position. Post-hoc pairwise testing was performed using Dunn’s test with a Bonferroni correction.

#### Classifying Wrist Posture from Extrinsic Finger Muscle EMG Activity

To investigate whether extrinsic finger muscle EMG signals might contain information about wrist posture, we developed a classifier to examine the capability of predicting wrist posture based on finger EMG activity alone. We trained two linear discriminant analysis classifiers, one for the extrinsic finger extensors and the other for the flexors, to predict if the movement of a digit was performed when the wrist was held in a flexed, neutral, or extended wrist posture. Each classifier was cross-validated using k-folds, with k = 5.

## Results

### Finger Movements at Different Wrist Postures

#### Finger Extensors

For all the finger extensor muscles, both posture and subject had a significant main effect on EMG activity (p<.0001, ANOVA). Further, the EMG activity of every finger extensor muscle was highest when the wrist was held in an extended posture. This activity was significantly higher than all other postures assessed (p<.001, Tukey’s HSD, **Fig. 3)**. In contrast, when the wrist was flexed, the finger extensors generated the lowest level of EMG activity during the extension phase of the motion, ranging from mean levels of 30 ± 13% to 53 ± 30% of their maximum activity level with the wrist was extended. This strong pattern of EMG activity in response to wrist posture was consistent across all of the extensor muscles with the highest levels of EMG activity occurring when the wrist was extended and the lowest levels occurring when the wrist was flexed. EMG activity for the finger extensors with the wrist in the neutral, pronated, and supinated postures was at an intermediate level and varied depending on the muscle. When the wrist was held neutral during digit extension movements, average EMG activity ranged from 47 ± 22% to 61 ± 35% of its maximum value with the lowest activity occurring in the D2 extensors (ED2, EIP) and progressively increasing for ED4 and ED5. Average EMG activity when the wrist was pronated ranged from 50 ± 28% to 67 ± 21%. Notably, the two index finger extensor muscles were differentially modulated when the wrist was pronated. While the ED2 and EIP activity was similar when the wrist was in the neutral, extended or supinated postures, the average EMG activity for ED2 and EIP was 50 ± 28% and 61 ± 26%, respectively when the wrist was pronated. This effect may be driven by the differing origin and insertion points of the ED and EIP muscles, which are accentuated during wrist pronation. With the wrist supinated, EMG activity across all the finger extensors varied the least, with all muscles having average activity levels between 49 ± 33% and 59 ± 31% of their maximum.

**Figure 3.**
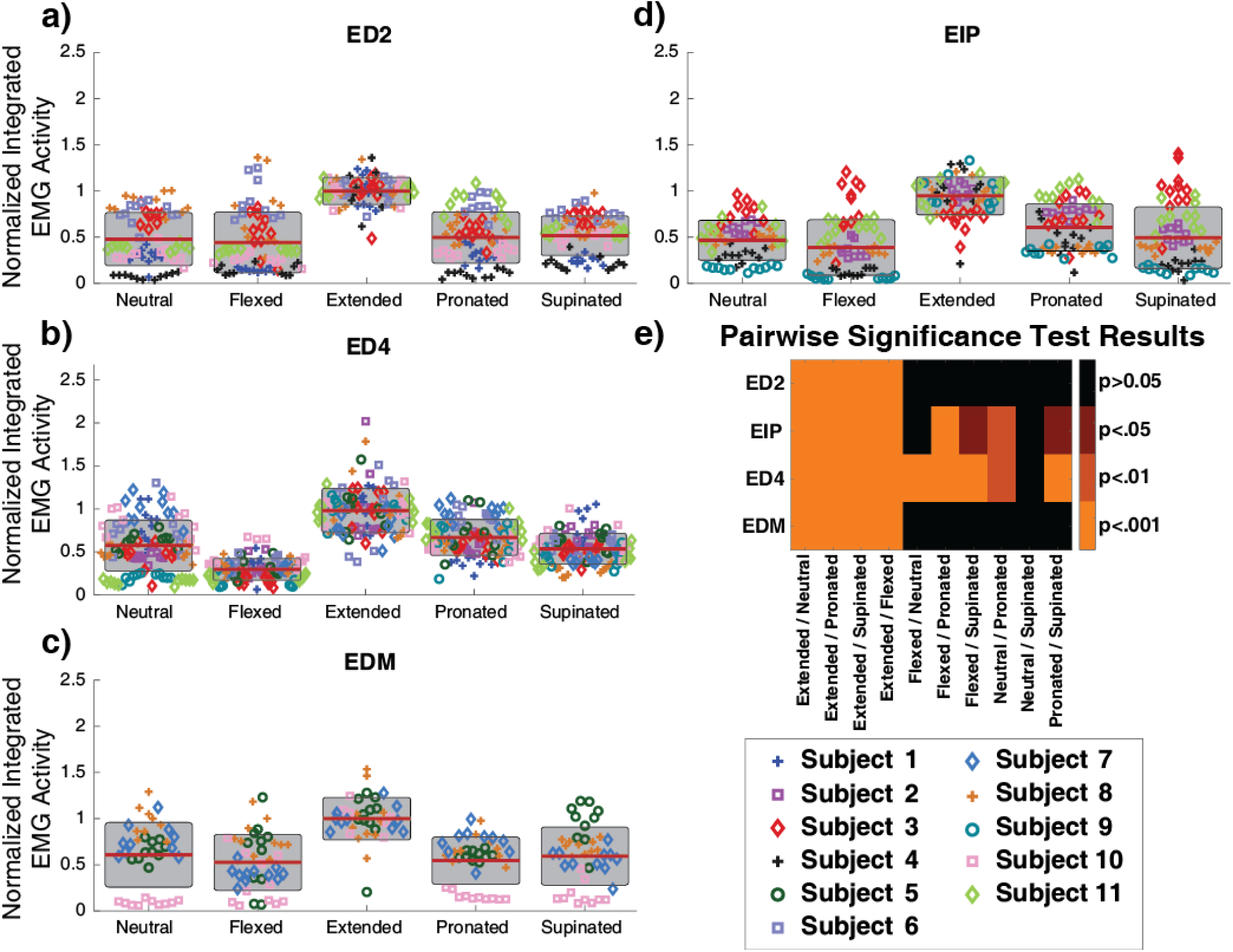
**(a-d)** Repetition and mean data of normalized integrated EMG activity for the extrinsic finger extensors performing extension actions during neutral, flexed, extended, pronated, and supinated postures. All finger extensors examined show increased EMG activity during finger extension movements when the hand was held extended. The horizontal red bars show normalized group means, and the standard deviation is shown as a gray box. Pairwise comparison of EMG activity of the finger extensors between the wrist extended and all other postures showed significant differences (p<.001, Tukey’s HSD). Individual data points represent each movement repetition and are color and marker coded for each subject across postures and plots. **e)** Heat map of pairwise significant differences for EMG activity of the extrinsic finger extensor muscle for all combinations of hand postures.

Of the extensor muscles assessed, ED4 showed the greatest difference in activity based on wrist posture. Of the ten possible pairwise comparisons (e.g. neutral vs. flexed, neutral vs. extended), all but neutral vs. supinated postures showed significant differences (p<.01, Tukey’s HSD). ED2 and EDM were affected the least by changing wrist posture; only pairwise comparisons testing extension against the other postures showed statistical significance **(Fig. 3e)**.

#### Finger Flexors

Similar to the finger extensors, wrist posture had a significant effect on the EMG activity of all of the finger flexor muscles (p<.001 for all muscles except FDS4, p<.05, ANOVA with Tukey’s HSD). The finger flexor muscles were most active when the wrist was held in the flexed posture, with the exception of FDS4 and FDP4, which were most active in a neutral posture with EMG activities of 87 ± 25% and 95 ± 26%, respectively. With the wrist in the neutral posture, the other flexor muscles had average activity levels that were much lower and ranged between 54 ± 18% to 73 ± 22% (**Fig. 4)**. For FDS4 and FDP4, the mean EMG activity when the wrist was flexed was 86 ± 29% and 87 ± 36% respectively, while the other finger flexors ranged from 86 ± 33% to 100 ± 22%. This result was unusual compared to the typical behavior of the finger flexors and may be due to anatomical differences in how the muscle lengths are affected by wrist posture. In contrast to the finger extensors, the individual finger flexor muscles showed little difference between extended, pronated, and supinated postures. Interestingly, the overall effect of wrist posture on EMG activity appeared to decrease for digits further away from D2 (**Fig. 4**).

**Figure 4.**
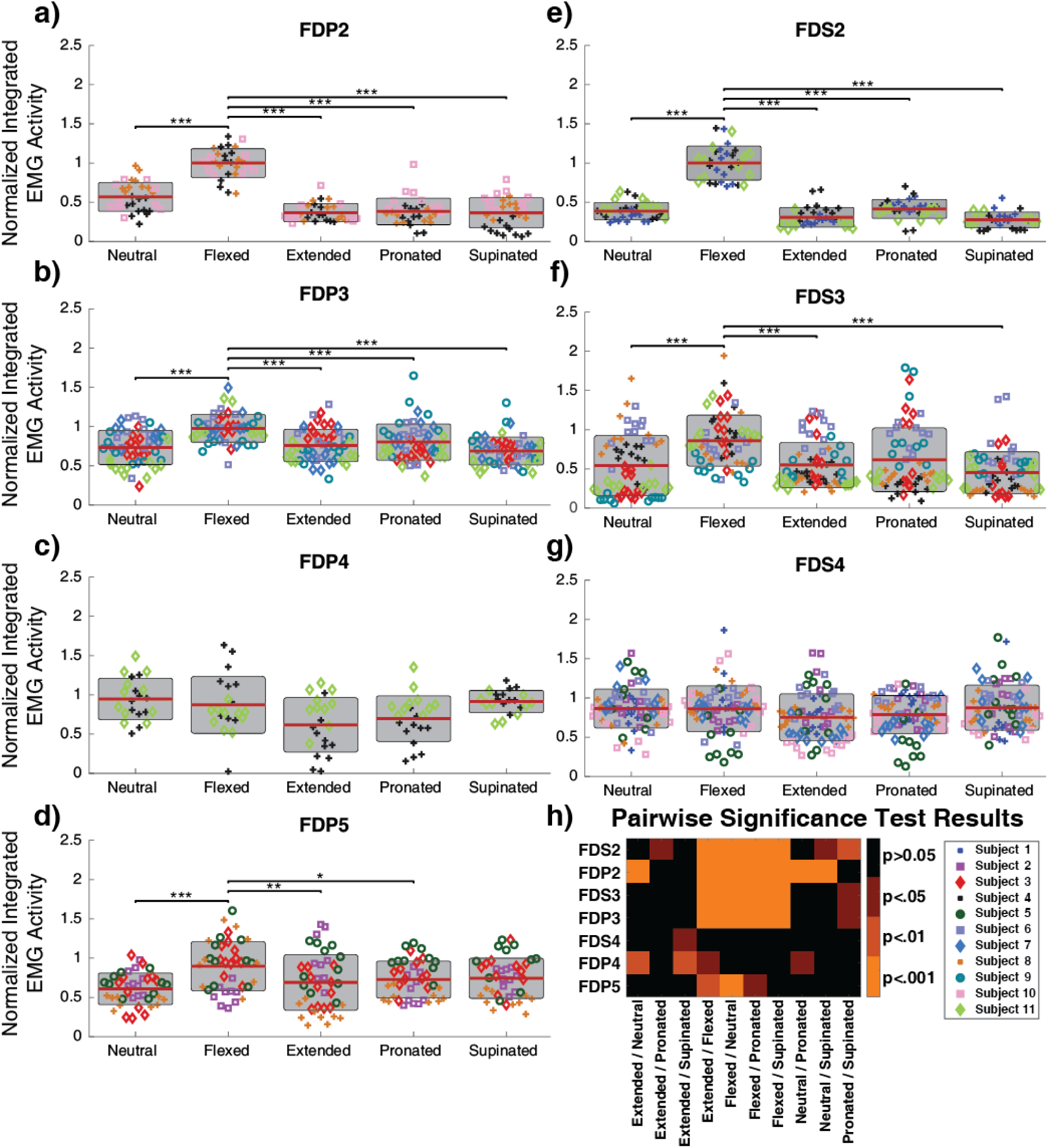
**(a-g)** Repetition and mean data of normalized integrated EMG for the extrinsic finger flexors performing flexion actions during neutral, flexed, extended, pronated, and supinated postures. The D2, D3, and D5 finger flexors showed significantly elevated EMG activity when the wrist was flexed. Individual data points represent each movement repetition and are color and marker coded for each subject across postures and plots. The horizontal red bars show normalized group means and the standard deviation is shown as a gray box. Significant differences between the flexed and other postures are denoted: * p<.05, **p<.01, ***p<.001 above the black bars (Tukey’s HSD). Further differences are shown in Table 4. **h)** Heat map of pairwise significant differences (Tukey’s HSD) of EMG activity between postures for the extrinsic finger flexors.

#### Inter-Subject Variance

For all muscles that were assessed, excluding FDS2, the individual subject had a significant effect on how wrist posture effected extrinsic finger muscle EMG activity (p<.05, Tukey’s HSD). This main effect response was primarily driven by high responders, which were subjects that had high levels of finger EMG activity when the wrist posture matched the action of the finger muscle (e.g. wrist extension for finger extensors). Subject 4 in **Fig. 3a** is an example of this. Similar to the other ED2 muscles, Subject 4 showed the highest level of activity when the wrist was held in the extended posture. However, unlike the other subjects, ED2 was much less active in all other wrist postures. Other examples of this phenomenon are EIP in Subject 9 (**Fig. 3d**) and EDM in Subject 10 (**Fig. 3c**). This effect was less pronounced in the finger flexor muscles.

### Extrinsic Muscle Finger EMG Activity During Wrist Movements

The extrinsic finger extensor muscles were highly active during wrist extension regardless of the finger posture. In fact, finger extensor EMG activity was significantly higher when the fingers were fully flexed and the wrist was being extended than when the fingers were extended and the wrist was being flexed (p<.001, Dunn’s, **Fig. 5a**). In the four combinations assessed (wrist extension/flexion with fingers extended/flexed), the extrinsic finger extensor EMG showed a wide range of activity levels. The highest EMG activity occurred during wrist extension when the fingers were held extended where the median normalized EMG activity from the finger extensors was 98% (IQR 85%-111%). When the wrist was extending and the fingers were held flexed, the median normalized EMG for the finger extensors declined to 52% (IQR 35%-77%). During wrist flexion movements, EMG activity was the lowest for the finger extensor muscles. When the fingers were held extended, median normalized EMG activity was 28% (IQR 16%-44%) and was even smaller at 18% (IQR 9%-33%) when the fingers were held flexed. The results showed significant differences in EMG activity for all pairwise comparisons of movement/posture combinations (p<.001, Kruskal-Wallis test with Dunn’s post-hoc test). Overall, we found that the finger extensor EMG activity changed by more than a factor of three when the fingers were held extended based on whether the wrist was moving in extension or flexion.

**Figure 5.**
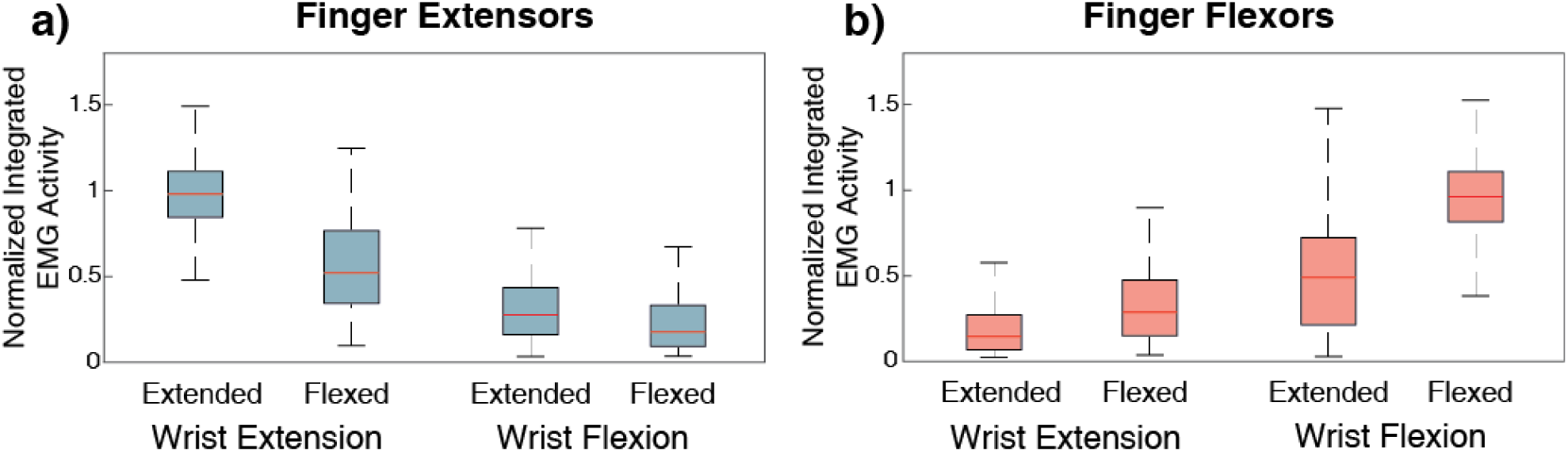
Normalized integrated EMG activity for **a)** finger extensors (N=22) and **b)** finger flexors (N=28) during wrist extension and flexion movements in flat and fist postures. For each subject, 10 repetitions were performed for movements in each posture. The red line shows the median, and the outer boxes are the first and third quartile. Error bars represent the 5-95% confidence interval. All posture and movement combination showed significant differences (p<.005, Kruskal-Wallis with Dunn post-hoc testing).

The EMG activation patterns of the finger flexor muscles showed a similar pattern to that observed in the finger extensors, with finger flexors significantly more active during wrist flexion movements than wrist extension movements, even when extension was performed with the fingers in a fist (**Fig. 5b)**. The highest EMG activity of the movements and postures assessed occurred during wrist flexion when the fingers were fully flexed at a median normalized EMG activity of 96% (IQR 82%-111%). The next highest level of EMG activity was during wrist flexion with the fingers extended with a median normalized value of 49% (IQR 21%-72%), nearly half of the highest value. Even when the fingers were being actively held in extension, finger flexor muscles were active during wrist flexion movements. EMG activity for the finger flexors was lowest during wrist extension movements; during extension movements with the fingers flexed, the median normalized EMG activity was 29% (IQR 15%-47%). When the wrist was extended and the fingers were extended, median normalized EMG activity was the lowest at 15% (IQR 7%-27%). The difference in activity levels were all significant (p<.001, Kruskal-Wallis test with Dunn’s post-hoc test) and varied by more than threefold depending on whether the wrist was flexing or extending when the fingers were fully flexed.

### Classifying Wrist Posture from Extrinsic Finger Muscle EMG Activity

Since we found that that extrinsic finger muscle EMG activity was often highly modulated by wrist posture, we asked whether wrist posture itself could be classified simply from the flexor or extensor finger muscle EMG activity. Overall, using just finger extensor muscles, the classifier accuracy was 63.5% ± 4.0% (99% confidence interval) and using just finger flexor muscles, the classifier accuracy was 47.4% ± 4.3% (99% confidence interval), both which were significantly above the chance level of 33%. True class prediction accuracy varied depending on the posture but followed the same pattern for both classifiers (**Fig. 6**). The most accurate true class predictions occurred when the wrist was held in the same posture as the finger action: wrist extended for the finger extensors (86.9%) and wrist flexed for the finger flexors (66.4%). When the wrist was in a neutral posture, true class accuracies were lowest for both classifiers at 32.7% for the finger extensors and 22.5% for the finger flexors. The confusion matrices and true class accuracy for both classifiers are shown in **Fig. 6**, and additional information on precision and recall is provided in **Supplementary Table 3**.

**Figure 6.**
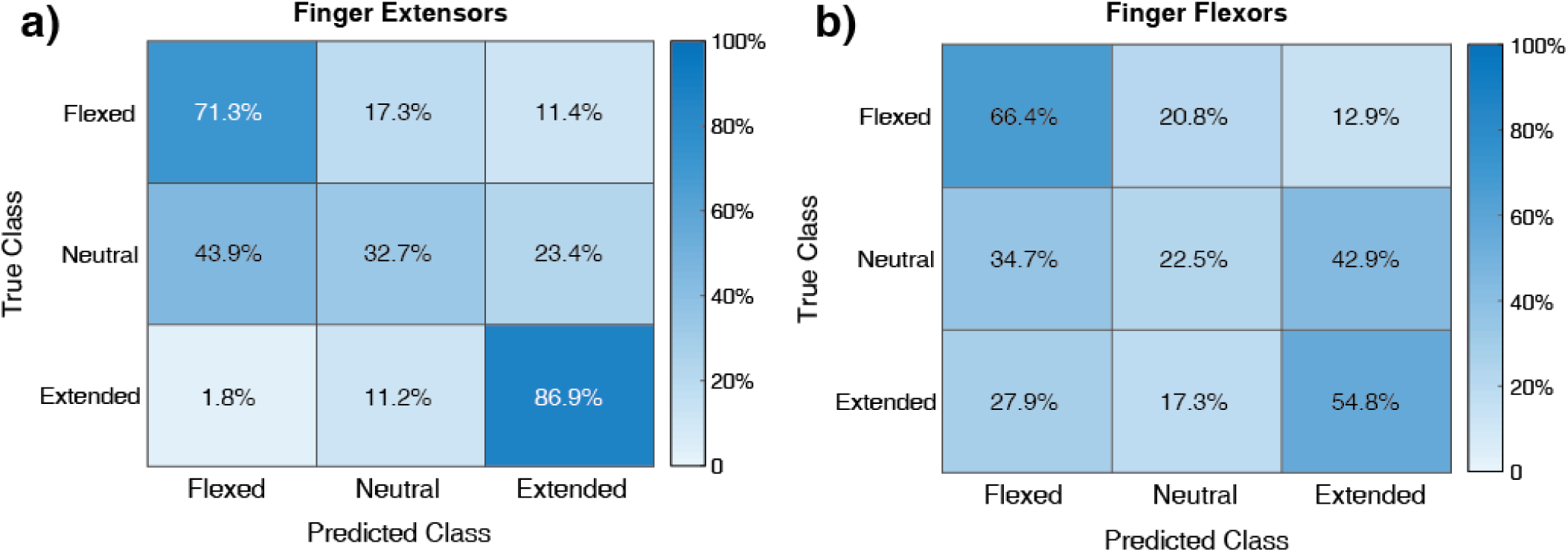
Confusion matrices and true class accuracy for the linear discriminant classifiers for the **a)** finger extensors and **b)** finger flexors. The diagonal of the matrices shows the percent of correct predictions (true positive rate); the other percentages in each row are the incorrect predictions of other hand postures (false positives). **a)** The finger extensors show the highest accuracy (86.9%) for detection of the extended wrist posture, and the lowest accuracy (32.7%) for neutral posture. The most common misclassification for the neutral posture was for the wrist flexed posture. For the finger extensors, the differences in EMG activity for flexed and neutral postures were minor and only significant for ED4, which may be the driving cause of this error. **b**) The finger flexors show similar results as the extensors, but with lower overall accuracy. Prediction of the flexed posture was the highest at 66.4%, followed by extended posture at 54.8%. Neutral posture classification had the lowest accuracy at 22.5%.

## Discussion

In this study, we found that there were wrist posture-dependent effects on extrinsic finger muscle EMG activity, which is consistent with previous reports^8,24,25,31^. We also expanded on this work by investigating EMG activity from the individual muscle compartments of the FDS, FDP and ED muscles, as well as using low-force movements that have not been previously studied. An important issue is that using intramuscular electrodes instead of surface electrodes increased the likelihood that the recorded signals were indeed from the same muscle in all postures. With surface EMG, there can be significant motion of the muscles under the skin, making it difficult to ensure that recordings at all wrist postures would be from the same muscle^8,24^.

In these experiments, we found that during single digit motions, both the finger flexors and extensors exhibited significant changes in their EMG activity based on the posture of the wrist, and in some cases varied by as much as 70% of the normalized value. We have also shown evidence that the extrinsic finger muscles have higher levels of activity during wrist movements that are performed in the same direction of action regardless of finger posture, which suggests that these muscles assist with unloaded wrist movements. These muscles may therefore also play a role in maintaining static wrist postures.

Posture dependent activation levels may be attributed to the altered tension of the extrinsic finger muscles caused by lengthening of muscles and tendons during different wrist postures, which change the force necessary to make such movements. The increased passive force of the extrinsic finger muscles when the is wrist extended and flexed has been previously documented^22,32,33^. For example, when the wrist is extended, the FDS and FDP muscles are lengthened, which generates an increased flexion torque on the digits. The extensor digitorum and other assisting muscles must therefore generate more force, and thus EMG activity, to counteract the increased tensions of the flexor muscles both to maintain finger position and to perform extension movements. During finger flexion movements while the wrist is still held extended, the lengthened finger flexor muscles generate substantial passive forces that assist in flexion movements and therefore require less EMG activity to make a flexion movement. This may also explain why the flexor compartments of the radial digits (FDS2, FDS3, FDP3) had the lowest activation in the supinated posture, while the flexor compartments of the ulnar digits (FDS4, FDP4, FDP5) were substantially more active in the same posture.

Despite the muscle compartments showing consistent patterns of responses, such as elevated EMG activity of the finger extensors when the wrist was extended and lower activity when the wrist was flexed **(Fig. 3)**, there were differences in the magnitude of those responses across fingers. This may be explained in part by the differences in passive forces generated by the individual compartments during hand postures. The study by Keir *et al*.^22^ showed that passive muscle force differences of up to 200% may occur between compartments of the same muscle, such as in the case of FDS2 and FDS4. It is therefore plausible that the extrinsic finger muscle compartments exhibit some differences in response magnitude based on wrist posture, but that the overall pattern of responses is maintained.

The accuracy of the classifiers used to determine wrist posture based on the integrated finger EMG activity were significantly above chance, demonstrating that information regarding wrist posture can be obtained from the extrinsic finger muscles. The high level of classifier accuracy for wrist postures in the agonist direction (wrist extended for the finger extensors, flexed for the finger flexors) suggests that information on wrist state can be gathered from the extrinsic finger muscles. However, information is limited to this preferred direction, due to the poor performance in predicting neutral posture. The performance of the classifiers is a reflection of the results reported in **Fig. 3** and **Fig. 4**; all muscles assessed showed at least some overlap between EMG activity in the neutral posture and the posture in the opposite direction of the action (flexed for the finger extensors, extended for the finger flexors), which caused difficulty in differentiating between these postures. Of the 11 muscles assessed, only 3 showed significant differences between EMG activity occurring in the neutral and antagonist postures, making it difficult to effectively distinguish between those postures when all muscles are taken into consideration. We chose to use separate classifiers rather than a single classifier to highlight the strengths of the flexors and extensors for predicting wrist posture in the flexed and extended postures respectively.

The shortcomings of the classification may also be attributed to using EMG activity of only extensor or flexor muscle groups for prediction. Motor control of the hand requires the coordination of multiple muscles for movements and stabilization of other joints. This holds direct implications for prosthetic control algorithms and was experimentally shown by the increased performance of the regression models that Smith *et al*.^20,34^ used for prosthetic control when they switched from using agonist/antagonist muscle pairs for dual-site control to including all recorded muscles in every degree-of-freedom calculation. If the activity of more than just an agonist/antagonist pair is not taken into account, there may be potential challenges in performing movements that use multiple degrees of freedom. For example, the dual-site formula by Smith *et al*.^34^ with ED4 and FDS4 as inputs would see three drastically different movements performed when the wrist was held extended, neutral, or flexed. Depending on where the threshold was set, the predicted movement could be a quick burst or no movement at all despite user intent of making a minor adjustment.

EMG activity was subject-dependent and found to be significant for nearly all muscles examined due to high responders. Interestingly, the EMG activity patterns of high responders appeared to be limited to a single muscle. For example, EDM of subject 10 showed high EMG activity during the extended posture but low activity during the other postures (**Fig. 3c**). In the other muscles and movements of subject 10, the EMG activity showed a range of responses similar to other subjects (**Fig 3a-b, Fig 4a,f)**. Although high responders had less pairwise differences in EMG activity due to posture, their performance still demonstrates that EMG activity for movements is affected by wrist posture.

A core limitation of our work was that we were unable to determine how much the change in extrinsic finger EMG activity based on posture was due to the necessity of overcoming mechanical coupling effects, such as the changing passive forces of antagonist muscles lengthening, or a motor control strategy in which the fingers assist in maintaining wrist posture. We are therefore unable to directly ascribe whether this phenomenon is a reaction to passive forces or a method of control that uses multiple muscles for force generation. Electrode placement was also a significant hurdle. Certain muscles were more difficult to target due to anatomical constraints and we were ultimately unable to successfully implant an electrode in ED3 due to the location of the posterior antebrachial cutaneous nerve. Electrode placement in FDS4 and FDP4 also proved challenging. Four electrodes that were targeted to FDP4 penetrated too deeply and were believed to be located in the flexor carpi ulnaris muscle in post hoc experiment data analysis, as the electrodes showed no response to finger movement but were active during wrist flexion and adduction. We were unable to determine whether this was due to the small size of these muscles or electrode migration that may have occurred when subjects moved their hands before testing. Only two electrodes generated high quality data from these muscles. Another limitation on our work was the testing of pronation and supination postures. While we did test finger movements in these wrist postures, we did not explore activity during pronation and supination movements as thoroughly as Mogk *et al*.^25^ and it was difficult to draw meaningful conclusions on these postures, as there did not seem to be a consistent behavior across the muscles examined. Biomechanical models of the arm and hand may assist in understanding these postures, as the EMG activity of the extrinsic finger muscles may be impacted by the changing muscle lengths and moment arms which can be modeled in simulation software.

In this study we demonstrated that wrist posture significantly influences the necessary action of finger muscles as demonstrated by the large changes in EMG activity of the extrinsic finger muscles in able-bodied subjects. Indeed, in certain cases, a three-fold change in finger muscle activity was required to produce the same finger kinematics when the wrist was held in different postures. For the finger flexors, these effects were most pronounced for muscles controlling the index finger. Future work could look to investigate whether the effects of wrist posture on finger muscle activity are still present in traumatic transradial amputees where these mechanical influences are removed, which could clarify whether these interactions represent a learned feed-forward control strategy driven by the cortex, or might also include sensory-driven feedback components.

## Data Availability

The dataset is available from the corresponding author on reasonable request.

## Acknowledgments

The authors of this study would like to thank Tyler Simpson, David Weir and Bree Bigelow for technical, engineering and regulatory support. We would also like to especially thank all the study participants. Research was sponsored by the U.S. Army Research Office and U.S. Army Research Laboratory and was accomplished under Cooperative Agreement Number W911NF-07-2-0055. The views and conclusions contained in this document are those of the authors and should not be interpreted as representing the official policies, either expressed or implied, of the Army Research Office, Army Research Laboratory, or the U.S. Government. The U.S. Government is authorized to reproduce and distribute reprints for Government purposes notwithstanding any copyright notation hereon.

## Author Contributions

C.R.B., M.M., L.E.F., J.L.C., M.C.M., M.L.B., and R.A.G designed the study. C.R.B., M.M., L.E.F., J.L.C., M.C.M., M.L.B., and R.A.G. conducted the experiments. C.R.B. analyzed the data. All authors contributed to the interpretation of the results. C.R.B. wrote the paper with M.M. and R.A.G., and all authors provided critical reviews, edits, and approval for the final manuscript.

## Supplementary Tables

**Supplementary Table 1:** List of movements performed for each subject.

| **Trial #** | **Movement** | **Wrist Posture** | **Finger Posture** |
| --- | --- | --- | --- |
| 1 | D2 Flexion/Extension | Neutral | Extended |
| 2 | D3 Flexion/Extension | Neutral | Extended |
| 3 | D4 Flexion/Extension | Neutral | Extended |
| 4 | D5 Flexion/Extension | Neutral | Extended |
| 5 | D1 Flexion/Extension | Neutral | Extended |
| 6 | D1 Abduction/Adduction | Neutral | Extended |
| 7 | D2 Flexion/Extension | Flexed | Extended |
| 8 | D3 Flexion/Extension | Flexed | Extended |
| 9 | D4 Flexion/Extension | Flexed | Extended |
| 10 | D5 Flexion/Extension | Flexed | Extended |
| 11 | D1 Flexion/Extension | Flexed | Extended |
| 12 | D1 Abduction/Adduction | Flexed | Extended |
| 13 | D2 Flexion/Extension | Extended | Extended |
| 14 | D3 Flexion/Extension | Extended | Extended |
| 15 | D4 Flexion/Extension | Extended | Extended |
| 16 | D5 Flexion/Extension | Extended | Extended |
| 17 | D1 Flexion/Extension | Extended | Extended |
| 18 | D1 Abduction/Adduction | Extended | Extended |
| 19 | D2 Flexion/Extension | Pronated | Extended |
| 20 | D3 Flexion/Extension | Pronated | Extended |
| 21 | D4 Flexion/Extension | Pronated | Extended |
| 22 | D5 Flexion/Extension | Pronated | Extended |
| 23 | D1 Flexion/Extension | Pronated | Extended |
| 24 | D1 Abduction/Adduction | Pronated | Extended |
| 25 | D2 Flexion/Extension | Supinated | Extended |
| 26 | D3 Flexion/Extension | Supinated | Extended |
| 27 | D4 Flexion/Extension | Supinated | Extended |
| 28 | D5 Flexion/Extension | Supinated | Extended |
| 29 | D1 Flexion/Extension | Supinated | Extended |
| 30 | D1 Abduction/Adduction | Supinated | Extended |
| 31 | Wrist Flexion/Extension | Neutral | Flexed |
| 32 | Wrist Flexion/Extension | Pronated | Flexed |
| 33 | Wrist Flexion/Extension | Supinated | Flexed |
| 34 | Wrist Flexion/Extension | Neutral | Extended |
| 35 | Wrist Flexion/Extension | Pronated | Extended |
| 36 | Wrist Flexion/Extension | Supinated | Extended |
| 37 | Wrist Abduction/Adduction | Neutral | Flexed |
| 38 | Wrist Abduction/Adduction | Pronated | Flexed |
| 39 | Wrist Abduction/Adduction | Supinated | Flexed |
| 40 | Wrist Abduction/Adduction | Neutral | Extended |
| 41 | Wrist Abduction/Adduction | Pronated | Extended |
| 42 | Wrist Abduction/Adduction | Supinated | Extended |

**Supplementary Table 2:** List of muscle targets and total number of electrodes placed.

| Electrode Location | Number of placements |
| --- | --- |
| EIP | 6 |
| ED2 | 7 |
| ED4 | 10 |
| EDM | 4 |
| FDS2 | 3 |
| FDS3 | 6 |
| FDS4 | 7 |
| FDP2 | 3 |
| FDP3 | 5 |
| FDP4 | 2 |
| FDP5 | 4 |

**Supplementary Table 3:**
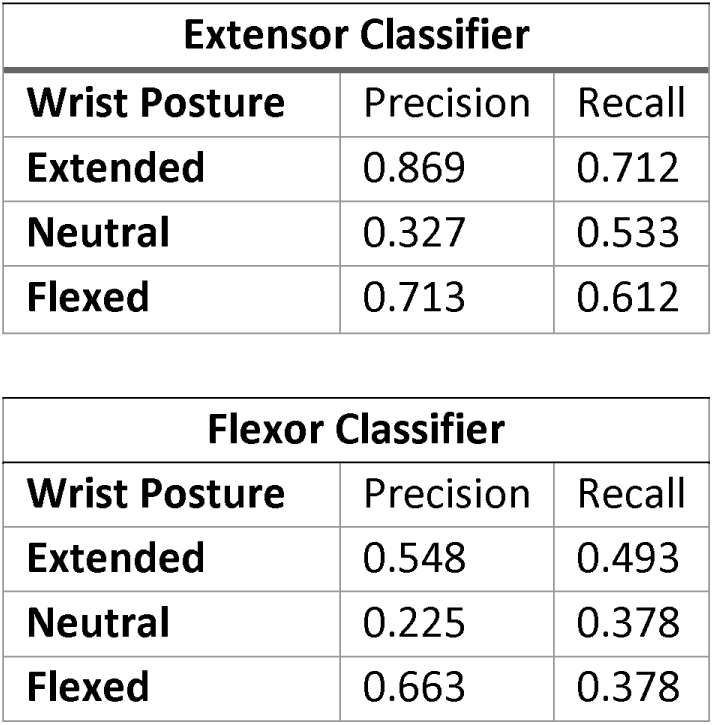
Precision and recall for posture classification using the extrinsic finger extensors (top) and flexors (bottom)

